# Structure-function analysis of Factor Seven Activating Protease (FSAP) by in-silico methods and its expression in insect cells

**DOI:** 10.1101/2025.01.07.631702

**Authors:** Sai Priya Sarma Kandanur, Abdulbaki Coban, Andreas Lange, Carsten Kemena, Aumkar Logendran, Bjørn Dalhus, Jonas Emsley, Sandip M. Kanse

**Affiliations:** Department of Molecular Medicine, Institute of Basic Medical Sciences, University of Oslo, Oslo, Norway; Institute for Evolutionary Biology, University of Muenster, Muenster, Germany; Department of Medical Biochemistry, Institute for Clinical Medicine, University of Oslo and Department for Microbiology, Oslo University Hospital, Oslo, Norway; School of Pharmacy, University of Nottingham, Nottingham, UK

**Keywords:** FSAP, HABP2, insect cells, recombinant, AlphaFold, structure, evolution

## Abstract

Factor VII Activating Protease (FSAP) is a circulating serine protease in blood and is likely to be involved in regulating hemostasis and inflammatory processes in stroke. The zymogen form of FSAP (Pro-FSAP) is activated by charged molecules, including heparin and histones. Here, we tested the feasibility of FSAP expression in insect cells and analyzed its structure and evolution with in-silico methods. The expression of full-length (FL) FSAP and the serine protease domain (SPD) was very low, and this was slightly improved with the protease-inactivating mutant. The remainder of the protein, comprising the heavy chain (HC), was expressed at high levels and demonstrated binding to pro-FSAP activators, histones, and heparin. Since FL-FSAP was not satisfactorily expressible for structural studies, the AlphaFold structure was used for molecular dynamics simulations and in-silico docking studies with pro-FSAP activators. AlphaFold predicts a globular structure for the EGF, kringle and the trypsin domain as well as a disordered N-terminal region (NTR). Docking studies suggested that the EGF-3 domain interacts with heparin and the NTR with histones. Very likely, this disturbs the globular fold of the enzyme to expose the activation site and promotes autocatalytic activation. FSAP evolved in its current domain configuration in jawed vertebrates about 500 million years ago suggesting an important and conserved function. The insights from these studies will facilitate the expression of recombinant FSAP as well as enable a deeper understanding of its structure and biological functions in the context of stroke.

## INTRODUCTION

Factor VII Activating Protease (FSAP) is a circulating serine protease that first evolved in the primitive jawless vertebrates (1). It has a broad substrate specificity and amongst other effects it, influences coagulation and fibrinolysis (2), stimulates protease-activated receptors (PARs) (3), increases vascular permeability in the lungs (4), degrades histones to reduce their toxicity (5–7), inhibits apoptosis (3) as well as releases nucleosomes from apoptotic cells (8). A polymorphism in the FSAP-encoding gene predisposes to stroke (9). Experiments in mouse models and *in vitro* studies (10) indicate that FSAP could be a potential treatment for ischemic stroke. However, inhibiting endogenous FSAP points to a more complex role in the context of stroke (11, 12).

Pro-FSAP is activated by polycations such as histones, protamine, and positively charged surfaces (6, 7, 13). Thus, the release of histones after tissue damage or inflammation, leads to the generation of active FSAP that degrades and detoxifies histones (5–7). This represents a reciprocal loop for the elimination of toxic histones from the circulation, analogous to the activation of the plasminogen pathway by fibrin to enable its own elimination (14). Pro-FSAP can also be activated by polyanions such as nucleic acids, heparin and dextran sulphate (15). FSAP has a high sequence homology to tissue plasminogen activator (tPA) and uPA, however, it is not a direct plasminogen activator (16). The domain organization of FSAP is very similar to that of the coagulation Factor XII (FXII) and the HGF family of growth factors (17). FXII (FXII) is activated by polyanions and results in the activation of coagulation (18). Thus, FXII and pro-FSAP represent canonical enzymes that are activated by charged polymers and may display mechanistic similarities in how zymogenicity is maintained and activation is initiated.

To date, the expression of recombinant full length (FL)-FSAP in HEK-293 and CHO cells was only possible in a restricted capacity, contingent upon removing its enzymatic activity (19, 20). FL-FSAP was also not expressible in *E. coli* (21). We reasoned that insect cells may be a more appropriate route as they have been used before for the expression of coagulation factors, Factor IX (22) and Factor XII (23). In this study, we have used a plasmid-based *Drosophila* expression system as well as the baculovirus expression system (24) to test the expression of FL-FSAP, SPD-FSAP and heavy-chain (HC)-FSAP in insect cells (Fig. 1A).

**Fig. 1:**
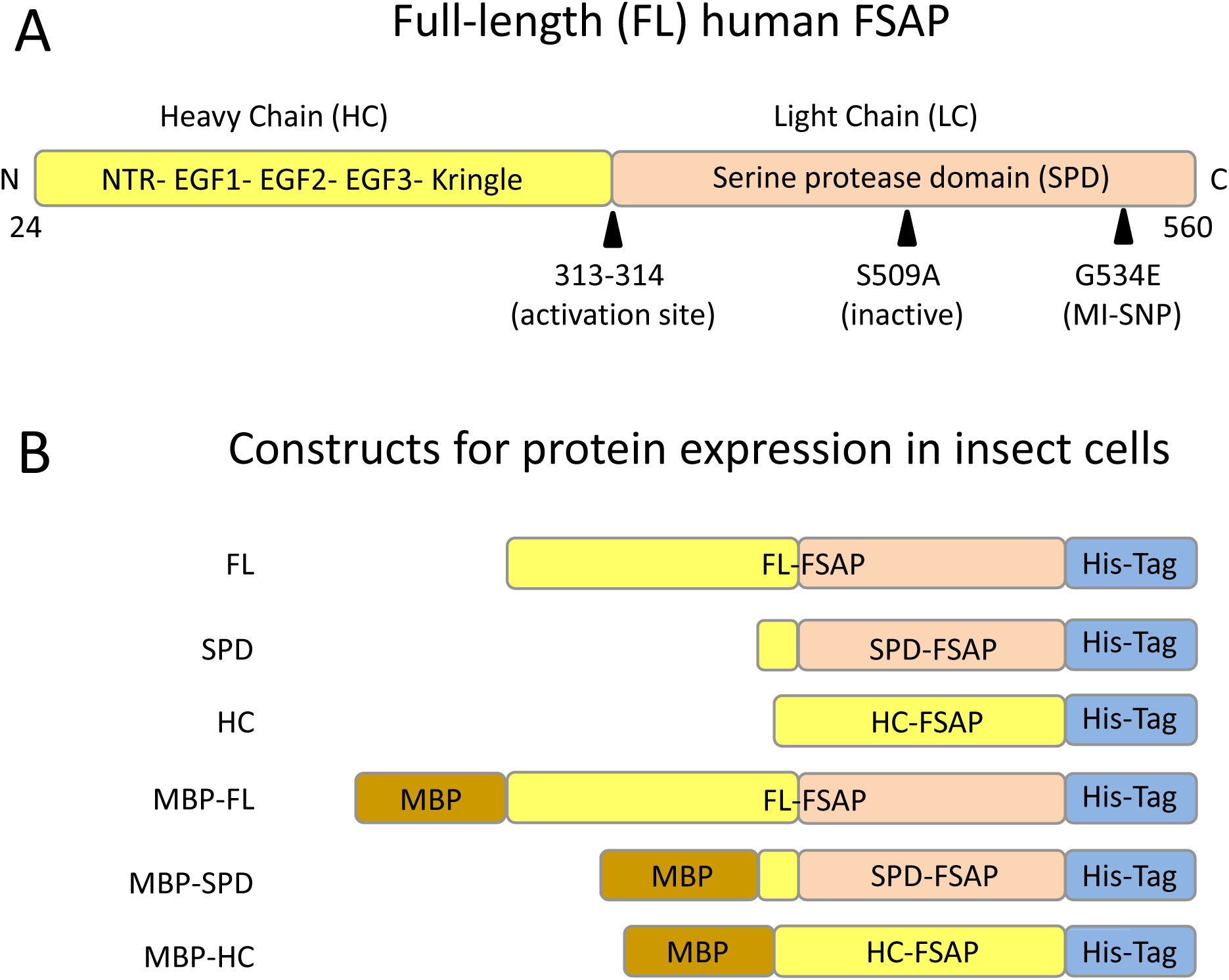
Constructs for the expression of FSAP in insect cells using the plasmid-based drosophila expression system: **A.** Schematic representation of full-length (FL) FSAP showing the serine protease domain (SPD) as well as the heavy chain (HC). The key amino acid positions are numbered according to the sequence of full length (FL) human FSAP. After cleavage of the signal peptide (1–23), single chain pro-FSAP consists of a 537 amino acid protein of which 24-72 represent the N-terminal region (NTR), 73-109 the EGF-1, 111-148 the EGF-2, 150-188 the EGF-3 domains, followed by 193-276 the kringle domain and 314-560 the serine protease domain (SPD). Activation occurs by proteolytic cleavage at position 313-314 (R/I) leading to the formation of a two-chain form that are connected by a disulphide bond. The SPD of FSAP is also denoted as the light chain (LC) (314–560) and the rest of the molecule constitutes the heavy chain (HC) (24–313). **B.** Constructs with and without codon optimization used in *Drosophila* expression system.

Until now, no structures of FSAP protein have been reported so we have analyzed its AlphaFold (25) model as well as performed molecular dynamic studies and docking analysis in-silico with potential zymogen activators, heparin (2) and histones (7). We have also investigated the evolutionary relationships of FSAP with other related proteases to better understand its structure and function. These studies lay the foundation for the analysis and interpretation of the structure, function, and the activation mechanism of FSAP.

## METHODS

### Plasmid-based *Drosophila* expression system (DES)

Plasmid (pMT-PURO), *Drosophila* Schneider *2* (S2) cells, culture medium and transfection reagents were obtained from ThermoFisher Scientific, Oslo, Norway. Full length (FL-) FSAP (gene *HABP2*) spanning residues 24-560, heavy-chain (HC-) FSAP (residues 24-291), and serine protease domain (SPD-) FSAP (residues 292-560) were cloned into pMT-PURO vector by Gibson cloning with a C-terminal 6X His-Tag (Fig. 1B). All constructs were also made in the vector pMT-PURO-MBP in frame with an N-terminal maltose binding protein (MBP) tag and a 6X His-Tag at the C-terminus for purification. A codon optimized form of human FSAP for expression in S2 was also used (GenScript, Piscataway, NJ).

S2 cells were cultured in Schneider’s *Drosophila* Medium supplemented with 10% fetal bovine serum (FBS) at 24°C and transfected using the calcium phosphate method. Transfected cells were selected with puromycin and adapted to serum-free Express Five (SFM) insect culture medium. Induction of expression was with CuSO_4_ (500 μM) under standard conditions. The effect of protease inhibitors e.g., aprotinin (Sigma, Oslo, Norway) (inhibitor of extracellular serine proteases) and BOS-318 (Selleckchem, Cologne, Germany) (inhibitor of intracellular furins) was evaluated as described before (26). Western blotting and ELISA were performed as described before (27).

### Purification of HC-FSAP

Conditioned medium (1 litre) was centrifuged to remove any cells and cell-debris and diluted 1:1 with buffer to obtain a final concentration of 20 mM HEPES (pH 7.4). This adjusts the pH and dilutes factors in medium that may interfere with Ni-NTA chromatography. After filtering through a 0.22 µm filter, the mixture was separated by Ni-NTA affinity chromatography (using a His-Excel column, Cytiva, Marlborough, MA) with equilibration buffer consisting of 20mM HEPES, 250 mM NaCl, pH 7.4, followed by a washing step with 10 mM imidazole and elution with a linear imidazole gradient (up to 1M). Relevant fractions were pooled, and buffer exchanged using Amicon^®^ Ultra -15 (Merck Millipore) with 10 mM HEPES, pH 7.4 to remove any salts from the sample and run on a HiTrap^TM^ Heparin HP 1ml column (Cytiva) with a linear NaCl gradient (up to 2M). As the final purification step, relevant fractions were pooled, concentrated, and run on a HiLoad^TM^ 16/600 Superdex^TM^ 75 pg column (Cytiva) in PBS and the resulting protein was concentrated.

### Dynamic Light scattering (DLS) of HC-FSAP

Purified HC-FSAP (1mg/ml) in PBS was tested for size and homogeneity of the solution. Samples were briefly centrifuged to remove aggregates and 10 µl was taken up in capillaries and placed in Prometheus Panta (NanoTemper Technologies, Munich, Germany) to measure its size and polydispersity Index (PDI). Samples were run in duplicates and data analysis was done using the software in the instrument.

### NanoDSF (Differential Scanning Fluorimetry) of HC-FSAP

The thermal stability of purified HC-FSAP was analyzed by nanoDSF (Prometheus NT.48, NanoTemper). 10 µl of 0.1 mg/ml HC-FSAP in PBS was taken up in capillaries and placed in the instrument. The temperature range for the measurement was 20-95°C, with gradual increase at 2°C/min. Samples were run in duplicate and the data was analyzed using the PR.ThermControl software (Version 2.3.1).

### Binding studies by MicroScale Thermophoresis (MST)

HC-FSAP (50 nM) was labelled using the Monolith protein labelling kit RED-NHS (NanoTemper) as per the manufacturer’s instructions. The labelled protein in PBS was added to a two-fold serial dilution of Heparin (Leo Pharma, Ballerup, Denmark) (starting at 6.6 µM), and Histone-H1 (Sigma) (starting at 66 µM), respectively. The samples were then measured at 20% LED/excitation power and medium MST power on Monolith NT.115 (NanoTemper). Data were analyzed using the MO Affinity Analysis software (version 2.3, NanoTemper).

### Alpha fold structure prediction, molecular dynamics and molecular docking

The predicted structure of Human-FSAP was obtained from the AlphaFold database (25) (https://alphafold.ebi.ac.uk/entry/Q14520). The electrostatic potential of the model was calculated using the APBS plugin in the PyMol Molecular Graphics System (version 2.4.0). The disorder prediction for full length FSAP was performed using DEPICTER2 (28), PSIPRED (version 4.0)(29) and IUPRED (version 3) (30). ClusPro 2.0 server (31) was used for complex prediction between AlphaFold pro-FSAP (AF-Q14520-F1) and heparin (tetra-saccharide model from the server itself) and Histone H3-H4 dimer (PDB ID: 7X57) with default parameters (32). All generated clusters were analyzed using PyMol.

To assess the stability of the predicted structures of FSAP (HC-, SPD-, and the FL-FSAP), we conducted molecular dynamics (MD) simulations using a modified version of a previously described protocol (33). The simulations were constructed and executed using the HTMD Python package (34). Each model system was solvated in an all-atom cubic box, with proteins centered at the origin of the simulation coordinates. We used water as the solvent and added NaCl ions to neutralize the system. Simulations were performed with the AMBER 14SB force field (35), starting with energy minimization and equilibration over 1 ns. Each system underwent a 200-ns simulation with the ACEMD engine (36) under default settings, in triplicate. Data analysis was carried out using HTMD and MDAnalysis (37).

For simulating the binding of heparin to FSAP, we utilized GROMACS 2022.1 (38). Structures were prepared following the standard GROMACS procedure. The system was solvated in a cubic box of SPC/E water with a 10-Å clearance, neutralized with sodium ions, then energy minimized and equilibrated. We ran three 200-ns simulations in an NPT ensemble, employing a V-rescale modified Berendsen thermostat at 300 K, a Parrinello– Rahman barostat at 1 atm, periodic boundary conditions, and particle mesh Ewald summation with a 1.6-Å grid spacing and fourth-order interpolation.

### Analysis of FSAP evolution

Twenty-four proteomes from ENSEMBLE (39), UNIPROT (40) and NCBI (41) databases were selected to represent most of the animal clades from Cnidaria to Chordata. Only the longest isoform was selected using ISOFORMCLEANER from DW-HELPER suite (https://zivgitlab.uni-muenster.de/domain-world/dw-helper). PFAMSCAN (42) was used to annotate protein domains using PFAM database v35 and INTERPROSCAN (v94.0)(43) to compare the annotation results. Reciprocal BLASTP (v2.12) (44) searches between human proteome and the 23 other species were run using ORTHOFINDER (v2.3.12) (45) and protein to genome EXONERATE (v2.4.0) (46) using human FSAP protein as query against genomes of the above mentioned species to be able to identify the origin of FSAP protein as accurately as possible. Domain-wise reciprocal BLASTP searches between FSAP domains from vertebrate species and the respective domains from basal chordates were performed to identify possible evolutionary histories as described before (47). MAFFT (v7.505) (48) was used to align the sequences and build phylogenetic trees using FASTTREE (v2.1.11) (49) with 1000 bootstrapping runs. SHOWTREE (https://github.com/LKremer/showTree) was used to visualize the phylogenetic tree with domain arrangements and TIMETREE (50) was used to construct the tree of FSAP with the evolutionary timescale.

## RESULTS

### Expression of FSAP in insect cells

In the baculovirus expression system (pFASTBac vector, Sf9/ Hi5 cells), low levels of expression of FL- and SPD-FSAP was observed in the intracellular compartment and none in the supernatant. Removing the His-Tag or replacing it with a FLAG-Tag; removing the honeybee melittin (HBM) signal peptide or replacing it with the FSAP signal peptide did not change this pattern and thus, this expression system was not used further (data not shown). Using the pMT-Puro plasmid-based Drosophila expression system, FL-FSAP and SPD-FSAP was not, or barely, expressed in S2 cells. However, HC-FSAP was expressed robustly in the cell extracts as well as in the conditioned medium (Fig. 2A). We then tested the effect of codon optimization on insect cell expression of FSAP. The pattern of expression of codon-optimized (CO) constructs was similar as that of the non-codon optimized (NCO) sequences across various experiments (Fig. 2). We also tested the effect of fusing MBP to the N-terminus of FSAP. None of the MBP fusion proteins were detected in either the cell extracts or the supernatants (Fig. 2). Thus, FL-FSAP and SPD-FSAP could not be satisfactorily expressed in insect cells, but HC-FSAP was expressed robustly using the DES.

**Fig. 2:**
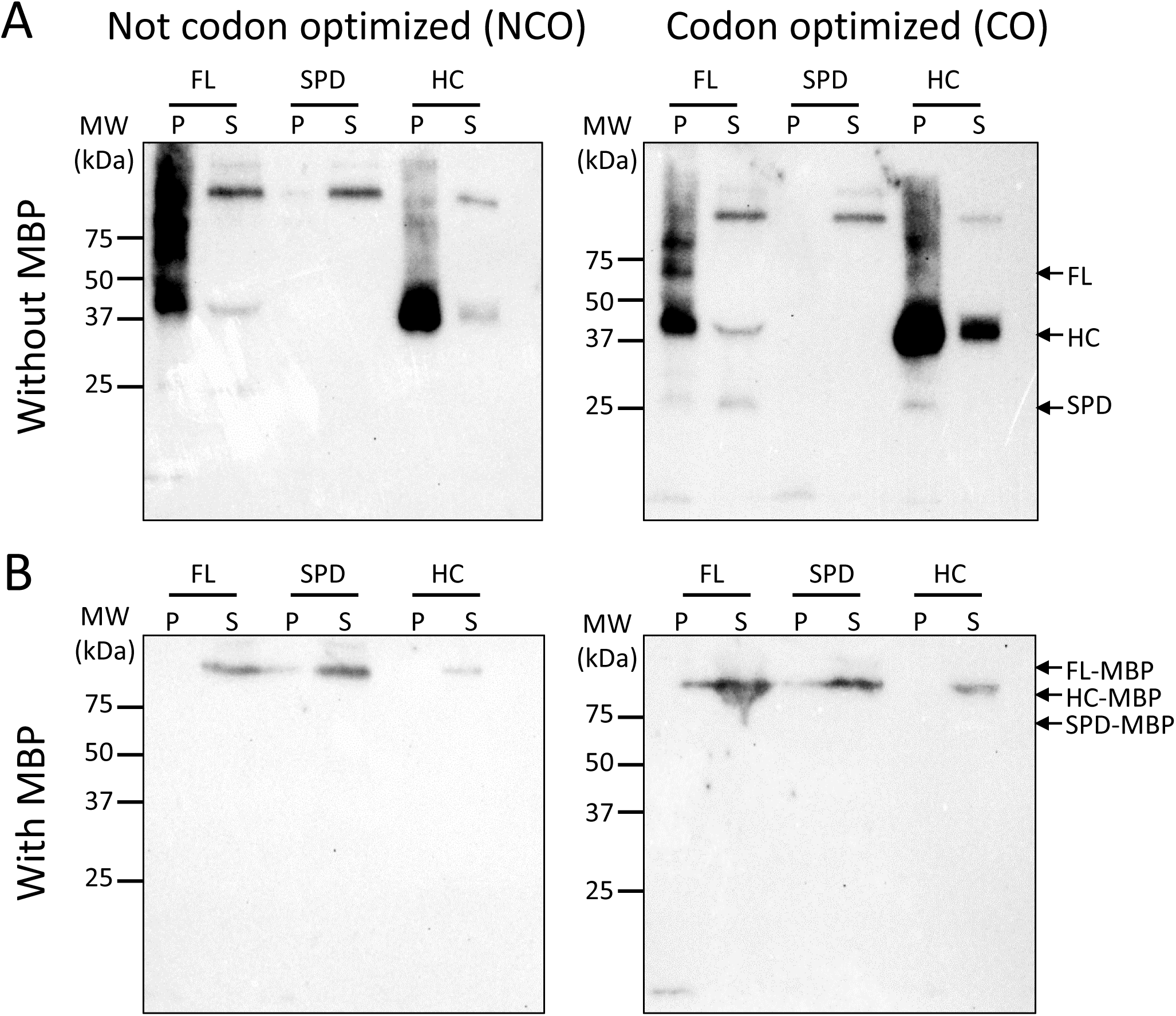
Expression of FSAP and its domains in insect cells: Western blot analysis of FSAP in the cell pellets (P) and supernatants (S) from S2 cells transfected with the different constructs. Expression without MBP **(A)** and with MBP **(B)**. Left panels are non-codon-optimized (NCO) and right panels are codon-optimized (CO). All samples are non-reduced and the expected MW of FL-FSAP is 65 kDa, HC-FSAP is 40 kDa, and SPD-FSAP is 25 kDa. MBP denotes maltose binding protein, and the MW of the fusion protein is expected to increase by 42 kDa. Similar results were obtained in three independent experiments.

### Effect of FSAP enzymatic activity on expression

Western blotting showed that the expressed FL-FSAP was often accompanied by bands corresponding to the SPD-as well as the HC-domain of FSAP indicating proteolytic cleavage inside the cells or in the supernatant. FSAP can cleave itself (51), but inhibition of FSAP by aprotinin did not alter this pattern (Supplementary Fig. S1). Intracellular furins play an important role in proteolytic processing of proteins in the secretory pathway (52) and their inhibition by BOS-318 did not inhibit this cleavage pattern (Supplementary Fig. S1). Mutating the active site of FSAP (S509A) increased the level of expression of FL-FSAP as well as SPD-FSAP (Supplementary Fig. S1) but the overall level of expression was very low when scaled up for protein purification (data not shown).

### Large-scale purification of HC-FSAP and its biophysical characterization

HC-FSAP was expressed at approximately 1 µg/ml in serum-free medium. It was eluted at ⁓800 mM of imidazole from the Ni-NTA column (Supplementary Fig. S2A), and at ⁓1.5 M NaCl from the heparin column (Supplementary Fig. S2B). HC-FSAP migrated at a MW of ⁓37 kDa in SDS-PAGE as expected, but the SEC chromatogram revealed that it eluted at a much higher MW (> 150 kDa) indicating that it adopts a multimeric or aggregated form (Supplementary Fig. S2C). Dynamic light scattering (DLS) analysis showed that the HC-FSAP solution is polydisperse with two broad peaks exhibiting a size of 4 ± 0.2 nm and 40 ± 77 nm, which also indicate substantial heterogeneity and multimerization (Supplementary Fig. S3A). A protein with 4 nm radius is estimated to be around 90-120 kDa large.

We also analyzed the thermal stability and unfolding of purified HC-FSAP by nanoDSF (Differential scanning fluorimetry). NanoDSF measures changes in the intrinsic fluorescence from tryptophan and tyrosine residues as the protein unfolds with increasing temperatures. Results indicate a folding state transition of HC-FSAP, where T_onset_ (the temperature at which unfolding begins) is 41°C and T_m_ (melting temperature) is 50°C (Supplementary Fig. S3B). Thus, HC-FSAP is uniform on SDS-PAGE but tends to form multimeric complexes.

### Binding of HC-FSAP to pro-FSAP activators

Anions like heparin (53), and polycations like histones (7) activate the zymogen form of FSAP and previous studies have indicated the involvement of HC-FSAP in this process (7, 27, 53). To this end, we analyzed the binding of purified HC-FSAP to unfractionated heparin, and Histone-H1 using Microscale thermophoresis (MST) (Fig. 3). The results show a mean *K_D_* value of 184 ± 86 nM and 2 ± 2 µM for HC-FSAP and heparin and Histone-H1, respectively. In MST experiments, conformational changes associated with ligand binding can either increase, as in Histone H1, or decrease fluorescence, as in heparin. Thus, HC-FSAP retains the binding to charged pro-FSAP activators. Both, unfractionated heparin as well as HC-FSAP are not well-defined monomeric molecules which will compromise the measured affinity values. ELISA-based binding studies were then used to further characterize this binding. A mixture of histones from bovine thymus, heparin-albumin conjugate, and bovine serum albumin (BSA) were immobilized. HC-FSAP bound strongly to histones but less so to heparin-albumin and not at all to BSA. The same pattern was observed with FL-FSAP purified from human plasma (Supplementary Fig. S4) indicating that HC-FSAP recapitulates the expected binding properties to heparin and histones. In additional binding studies, we found binding H3/H4 and H2A/H2B complexes to FSAP (data not shown).

**Fig. 3:**
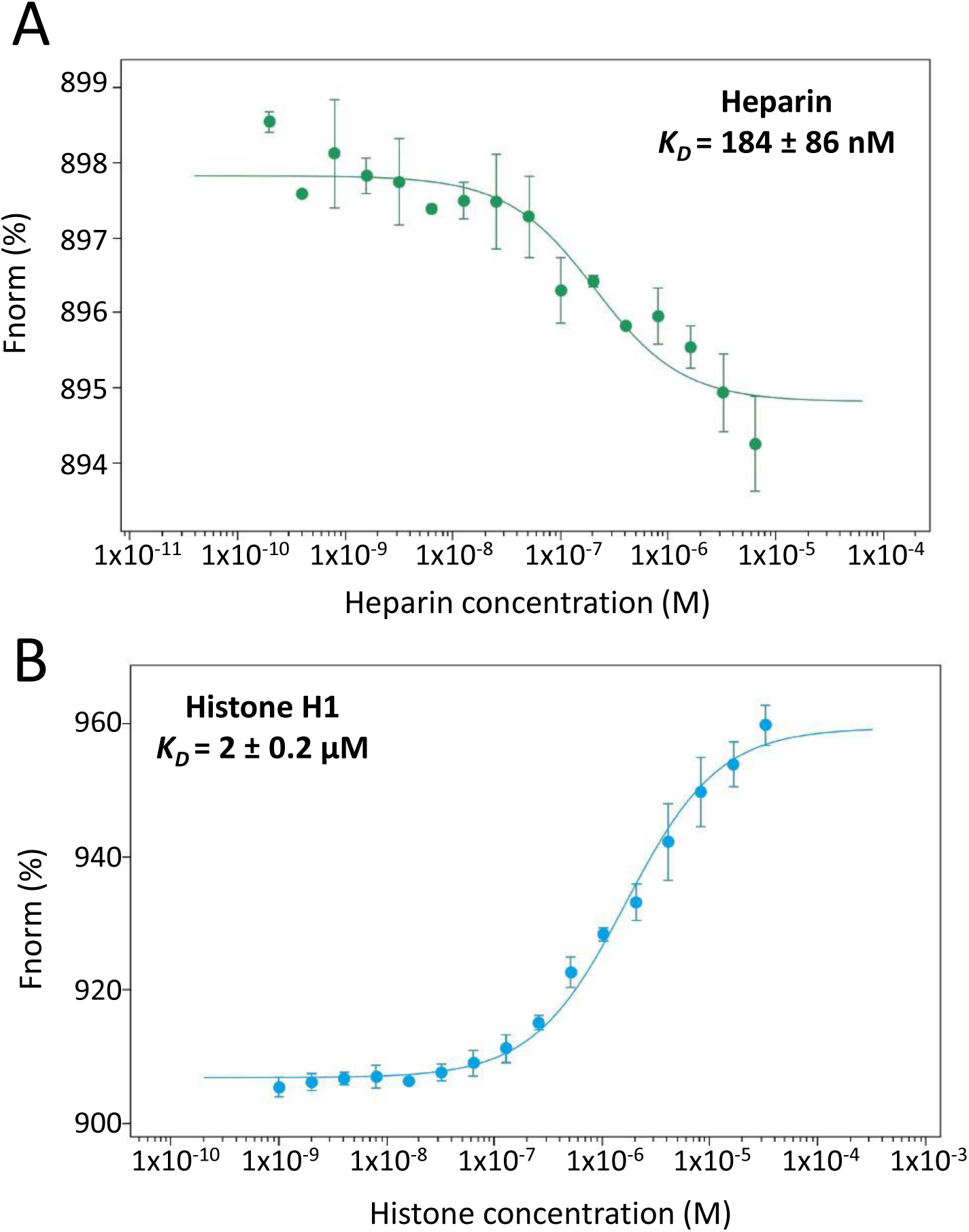
Microscale thermophoresis (MST) of HC-FSAP binding to its activators: Serial dilutions of **A.** Heparin (starting at 6.6 µM) and **B.** Histone-H1 (starting at 66 µM), were titrated against 50 nM of labelled HC-FSAP. The interaction between the molecules were measured using Monolith NT.115. The *K_D_* values from each analysis are depicted as mean ± SD, where n=3 independent experiments.

### 3D AlphaFold structure of pro-FSAP

AlphaFold (25) predicts the structure of the individual domains with high reliability (Fig. 4A, blue ribbons), but the NTR and the interconnecting loops are modelled with low confidence (Fig. 4A, orange ribbons) (54). Charge distribution analysis shows a cluster of basic residues on the surface of the EGF3 domain as well as a weaker cluster in the SPD domain (Fig. 4B). Acidic regions were present in the NTR, kringle domain-facing the activation site and most prominently around the active site in SPD. The active site as well as the activation site are buried inside the protein and the positioning of the loops in the SPD domain are in their zymogen form. (Fig. 4B). The predicted aligned error (PAE) plot predicts interaction between the EGF1, EGF2, EGF3 and the kringle domain (Supplementary Fig. S5) with low expected position error (between 0-5 Å). Interactions involving the other domains, NTR and SPD, were not predicted in this model as shown by the high positional errors.

**Fig. 4:**
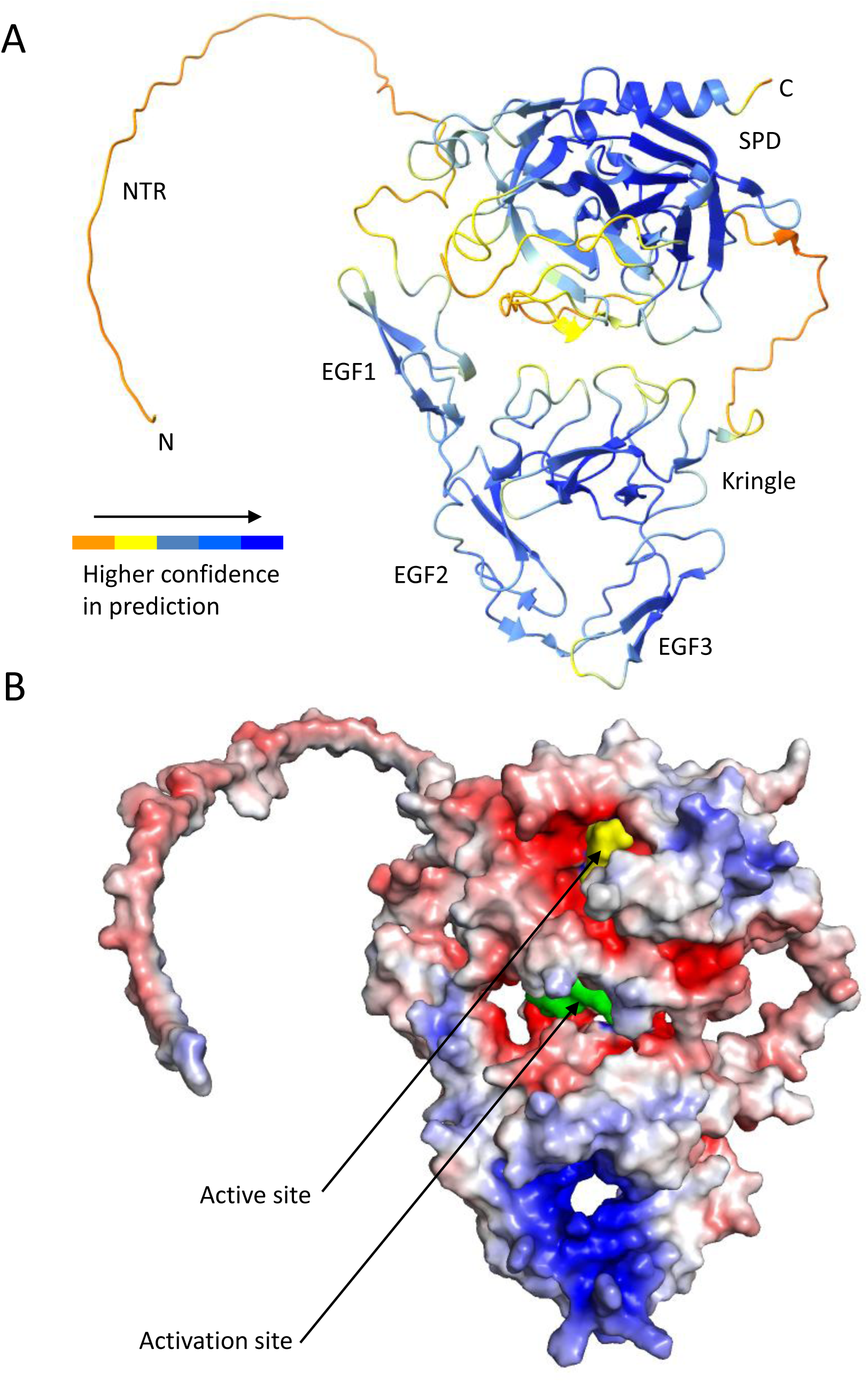
AlphaFold structure of FL pro-FSAP. **(24–560): A.** The predicted structure of pro-FSAP coloured according to the confidence score (pLDDT values) ranging from low (orange) to high (blue) https://alphafold.ebi.ac.uk/entry/Q14520. The domains of the protein are noted: NTR, EGF1, EGF2, EGF3, Kringle, SPD as well as the N-and the C-terminus. **B.** Surface charge distribution of basic residues (blue) and acidic residues (red) with the active site (yellow) and activation loop (green).

The primary sequence of full-length FSAP was analysed for intrinsic disorder using DEPICTER2 (28) which predicts disorder for residues 50-60 in the NTR. Other prediction servers, IUPRED and PSIPRED confirmed this. In addition, PSIPRED also identified disordered regions involved in protein binding within the central and C-terminal region of full-length FSAP (Supplementary Fig. S6). Thus, different analysis indicates that the NTR of FSAP is highly disordered.

### Pro-FSAP activation

The site of cleavage leading to pro-FSAP activation is buried at the center of the structure between the protease domain and the kringle/ EGF domains and this inaccessibility for interactions with cofactors or other proteases may be important for maintaining zymogenicity (Fig. 4B). It has been suggested that the EGF3-NTR interaction is important for maintaining the “closed” zymogen conformation and that activation is associated with the disruption of this interaction (55). However, in the AlphaFold model of FSAP zymogen, no proximity between EGF3 and NTR was predicted which might be related to the difficulty in assigning a specific location to the disordered NTR.

The interaction between heparin and the AlphaFold model of pro-FSAP was analyzed using ClusPro 2.0. In the best models, one molecule of FSAP interacted with two molecules of tetra-saccharide heparin. Results show interaction with heparin via residues Arg170, Arg171, and Lys173 through electrostatic interactions (Fig. 5A). Parameters related to ClusPro results are provided in Fig. 5C. The neighboring positively charged amino acids, Arg122, Lys139, and His140 of the EGF2 domain, and Lys185 and Arg208 may also play a role in stabilizing this interaction with heparin. This confirms the results of previous studies demonstrating that polyanions such as heparin bind to the positively charged amino acids in EGF3 domain to facilitate auto-activation of FSAP (53).

**Fig. 5:**
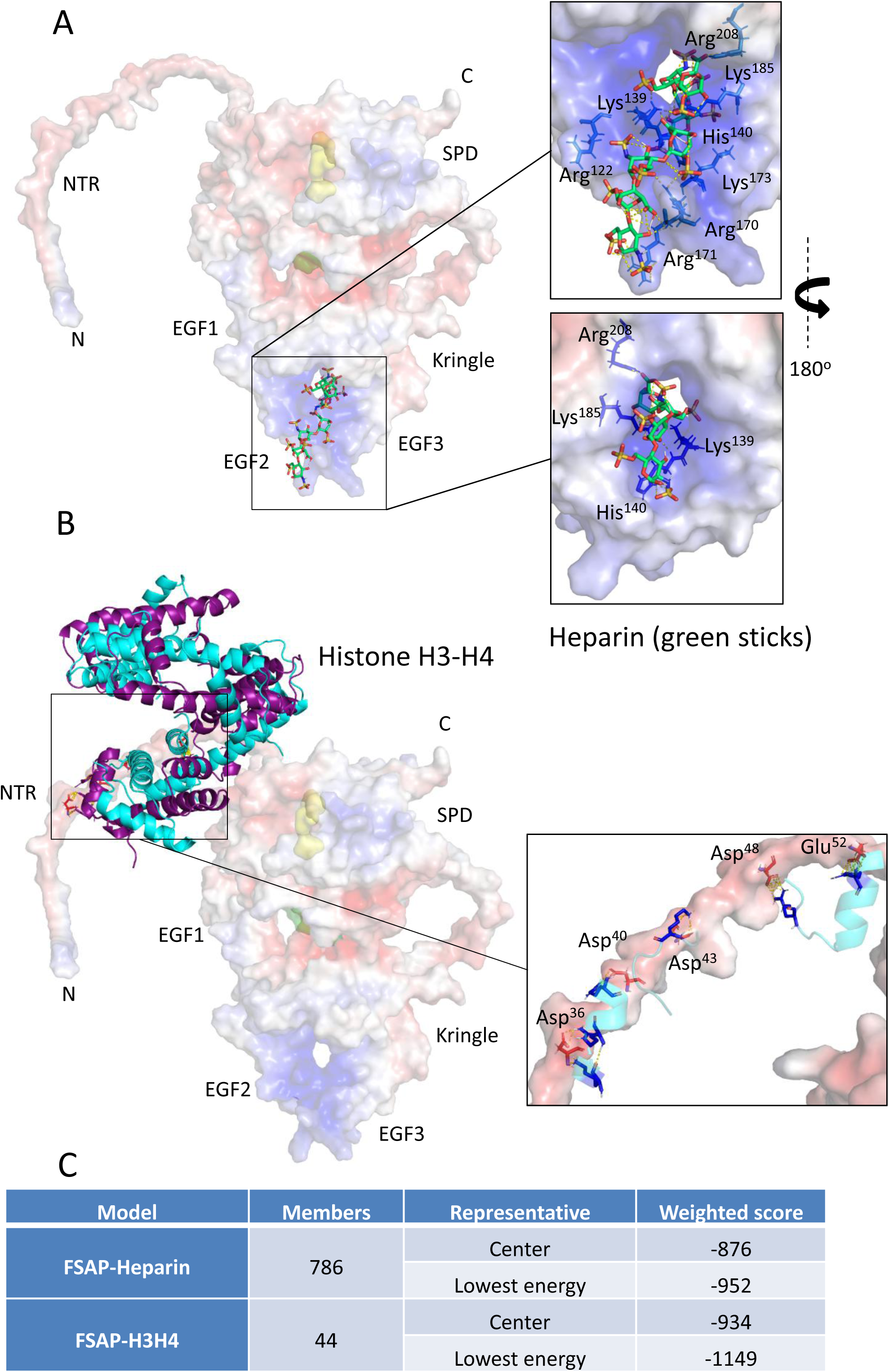
Putative interaction between FL pro-FSAP and its activators heparin and histone H3-H4: **A.** Model of pro-FSAP in complex with 2 molecules of heparin, each comprising of a tetra-saccharide, (green, ball and stick) and **B.** Histone H3 (cyan)-H4 (purple) dimer (PDB ID-7X57). FSAP is shown as a solid surface with its negative (red) and positive (blue) electrostatic potential, active site (yellow) and activation site (green) are depicted. Inset: The residues (stick model) involved in polar interaction with heparin and histones are zoomed in respectively. These complexes were predicted using ClusPro 2.0. For each docking analysis, the top 10 clusters generated upon docking were inspected for plausible interactions. One credible cluster was chosen and analysed for interaction sites. C. Members represent the cluster size. The weighted scores for the centre and the lowest energy of the cluster are indicated in the table.

Analysis of the interplay with histone H3-H4 (PDB ID: 7X57) suggests interactions with the negatively charged residues Asp36, Asp40, Asp43, Asp48, and Glu52 in the NTR of FSAP with a score of -934 (Fig. 5B). Thus, each of the activators of pro-FSAP appear to interact with a different region of the protein to initiate the autocatalytic activation process through the involvement of the NTR-EGF3 axis.

### Molecular Dynamic simulations

To refine the results obtained from AlphaFold, we conducted three independent MD simulations for (i) SPD, (ii) HC- and (iii) FL-FSAP using the HTMD Python package. In all three simulations, the SPD domain of the FSAP protein exhibited lower RMSD (root mean square difference) values, reduced fluctuations, and high compactness. Mapping the RMSF (root mean square fluctuation) values onto a representative structure of the SPD domain revealed consistent regions of conformational flexibility across each replicate. Areas of high flexibility are Pro336 to Gly344, Pro419 to Leu426, and Val454 to Glu463 (red areas Figure 6, left column). The HC and the FL protein displayed a highly flexible loop in the NTR of both structures (in HC-FSAP: Asp36 to Tyr46 and in FL-FSAP: Phe24 to Ser56, Fig. 6 middle and right column). Apart from the significant wobble in NTR, the rest of the structures remained consistent with the predicted AlphaFold models. The HC has one additional highly flexible area (Lys260 to Tyr270) and the FL-FSAP from Trp256 to Cys276.

**Fig. 6:**
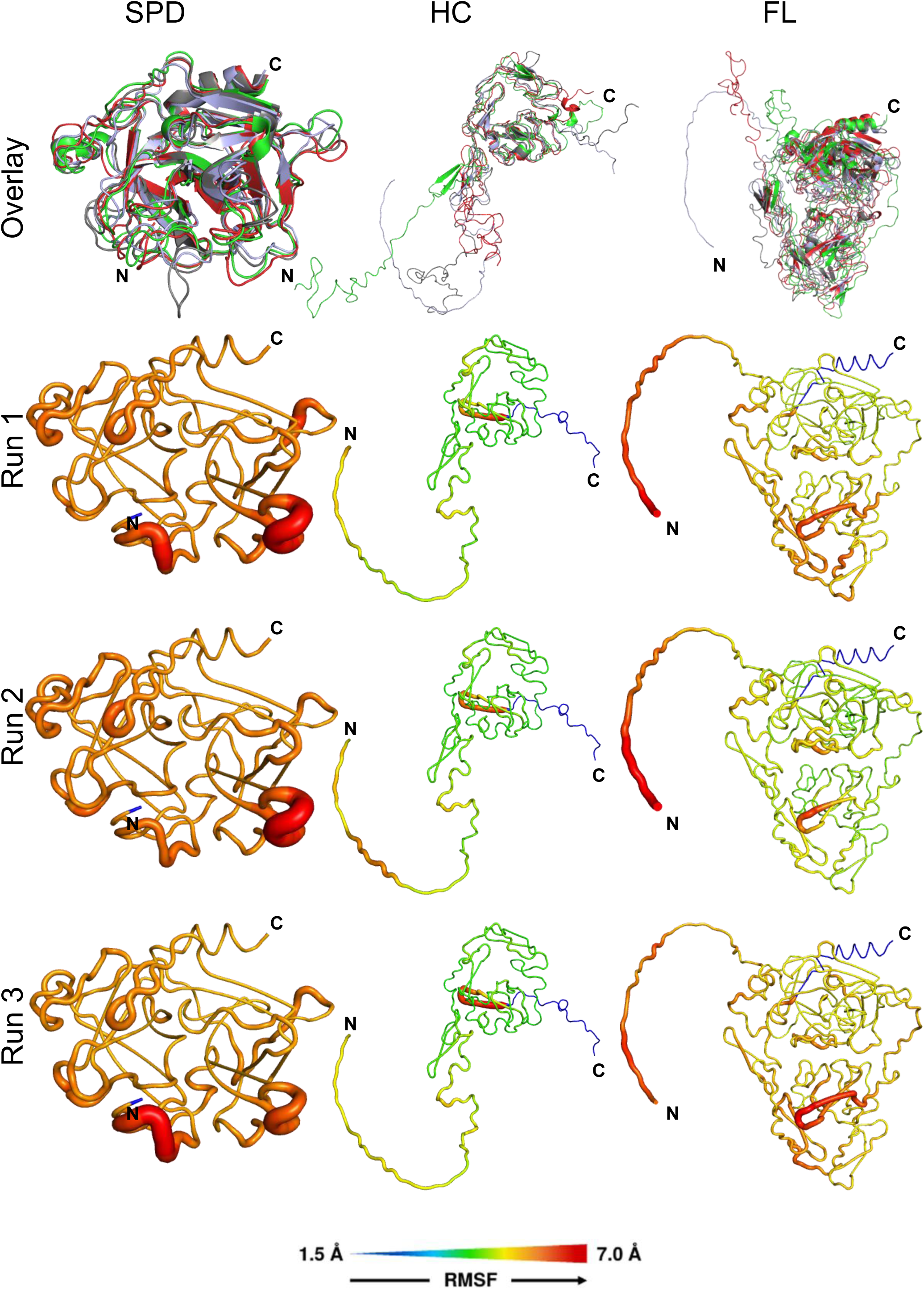
MD simulation of the SDP-, HC- and FL-FSAP: The first row displays an overlay of the initial AlphaFold-predicted structure (used as the starting structure for MD, shown in light blue) and the MD simulation results after 200 ns for each run: run1 (red), run2 (green), and run3 (gray). The rows labeled run1 to run3 show the RMSF values mapped onto a representative structure from each respective MD simulation. These mappings highlight similar regions of conformational flexibility across the replicates, where increased flexibility is indicated by more red (higher RMSF), and stability is indicated by dark blue (lower RMSF). This visualization allows for a clear comparison of structural stability and flexibility across the three simulations for each FSAP protein variant.

Additionally, we conducted MD simulations of the FL-FSAP protein bound to heparin using GROMACS. The Heparin-FSAP complex, as predicted by AlphaFold, remained highly stable and compact in all three independent runs, mirroring the stability observed in the individual protein structures. Two heparin molecules, each a tetra-saccharide, were predicted to bind to FSAP, so we ran three independent MD simulations for each molecule separately (Supplementary Fig. S7). In five out of the six simulations, the heparin molecules remained quite consistent with the predicted initial starting structures. In one simulation, however, one of the heparin molecules moved towards the position typically occupied by the other molecule. Despite this, it remained within the appropriate binding region, reinforcing the accuracy of the predicted heparin binding site. In summary, the MD simulations closely reflect the AlphaFold-predicted structures, with minor fluctuations.

### Evolution of FSAP

In the 24 studied species (Supplementary Fig. S8A), no orthologue(s) of FSAP protein in Cnidaria (*S. pistillata*) and Echinodermata (*S. purpuratus*) clades could be identified by EXONERATE, BLASTP and ORTHOFINDER searches. As for Cephalochordates and Tunicates, there were reciprocal BLASTP hits between human FSAP and *B. floridae* and *O. dioica*, but only partial hits using EXONERATE. The branches containing lower chordate candidates for FSAP in the constructed gene tree also have low bootstrap values (Supplementary Fig. S9). Besides, FSAP has a conserved domain arrangement in jawed vertebrates: EGF, EGF, EGF, Kringle, trypsin/SPD while the matched proteins have different domain arrangements in *B. floridae, O. dioica*, and *C. intestinalis* (Supplementary Fig. S8B).

ORTHOFINDER groups uPA, tPA and FSAP together in one orthogroup in the jawed vertebrates (Supplementary Fig. S9). In the lampreys there are two sequences and both cluster with FSAPs from other vertebrates, suggesting a possible gene duplication (Supplementary Fig. S8). Reciprocal BLASTP searches revealed that the kringle and trypsin domains of the protein found in Cephalochordates (XP 035669535) and Tunicates (ENSCINP00000001165) closely match those of vertebrate FSAP proteins, indicating a shared ancestry. However, the jawless vertebrates exhibit a distinct domain arrangement (Fig. 7). While one copy has an extra fn2 domain on the N-terminus, the other copy in *E. burgeri* has only one EGF domain instead of canonical three EGF domains. In the higher vertebrates, we identified orthologues of FSAP in every studied species including bichir species (*P. senegalus* NCBI protein ID: XP 039591852.1) in which it was previously reported to be missing. Considering the orthology analysis and the present domain arrangements in basal chordates, FSAP potentially emerged 600 million years ago (mya) (Fig. 7) in jawless vertebrates. About 500 mya, the addition of an EGF domain to the N-terminus created the jawed vertebrate FSAP, which has since then remained unchanged across all vertebrate clades (Fig. 7).

**Fig. 7:**
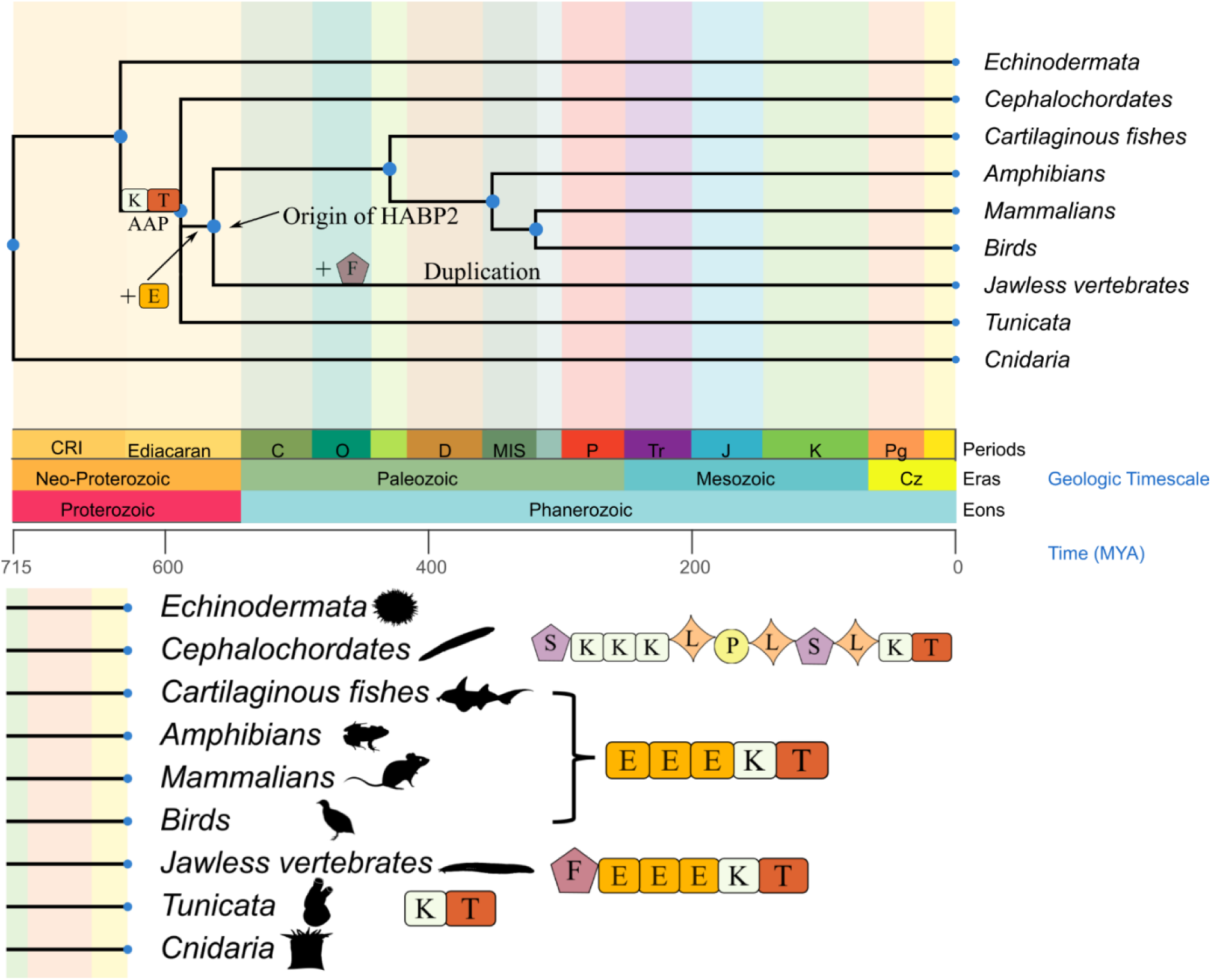
Evolution of FSAP (HABP2 gene): Step by step evolution of FSAP with the respective evolutionary timescale. Events on the domain level are marked on the tree. FSAP protein arose around ∼600 mya (million years ago) with the multiple EGF domain gains on the assumed ancestral protein. In jawless vertebrates, gain of fn2 domain and duplication event took place. The jawed vertebrate form of FSAP emerged ∼500 mya. AAP stands for “Assumed Ancestral Protein. Domain annotations: E: EGF, K: Kringle, T: Trypsin/SPD, F: fn2, S: SRCR, L: Ldl-a and P: PAN.

A conservation analysis of FSAP across 81 different vertebrate species from the phylum vertebrata (Supplementary Fig. S10) indicates a strong sequence homology within the defined structural domains such as the SPD, Kringle and EGF domains, while the NTR shows higher variability.

## DISCUSSION

Structure analysis needs large quantities of high-quality proteins which prompted us to test the utility of baculovirus-as well as plasmid-based expression system to express recombinant FSAP. In the baculovirus expression trials, the FL-FSAP protein was expressed in very low amounts in the intracellular compartment which recapitulated earlier results in mammalian cells (20). FL-expression in the DES system was also very low and this problem was not overcome by codon optimization, fusion with MBP or by using protease inhibitors. Mutation to inhibit the enzymatic activity of FSAP (S509A) improved expression but upscaling was not successful. Thus, over-expression of full-length human FSAP appears be toxic for insect cells.

HC-FSAP was expressed in insect cells at a high level which enabled successful purification and biochemical characterization using ELISA-based binding assays, DLS, nanoDSF, and MST. The results indicate that purified HC-FSAP exhibits reasonable binding affinity to its activators, heparin as well as histones and that it exists as a high MW species indicating multimerization/ aggregation. It is difficult to precisely identify which oligomeric forms are present due to the extended NTR which is likely to increase the hydrodynamic radius of the protein. To conclude, insect cells are well-suited to express HC-FSAP.

In the AlphaFold model, parts of the protein comprising of well-defined domains were predicted with high reliability (low PAE values), but the NTR as well as the interconnecting regions were predicted to be unstructured and flexible. It is known that even regions with highly reliable predictions can deviate to a large extent from experimental structures and thus the results should be interpreted with caution (56).

Activation of FSAP zymogen with positively and negatively charged molecules is akin to the activation of the intrinsic pathway of coagulation by negatively charged surfaces. The activation of FXII by negatively charged surfaces is attributed to clusters of positive residues on the surface of the heavy chain and this is confirmed by domain deletion and antibody inhibition studies (57). In FSAP, the positively charged EGF3 domain binds strongly to heparin in our models confirming earlier binding analysis (20, 53). FSAP zymogen activation by histones, by analogy, should target negatively charged regions. One such region is the NTR, but its intrinsic disordered nature excludes more detailed analysis in our model despite experimental evidence favoring this domain (27, 55). Intrinsically disordered regions/ proteins tend to be integrating foci of signaling pathways due to their flexibility and ability to undergo function-driven conformational transitions and the same may be the case for the NTR of FSAP (58). The disordered nature of NTR may also interfere with crystallization and cryo-electron microscopy studies on FSAP.

Proteases and their substrates can exhibit convergent or divergent evolution. The evolutionary analysis of FSAP sequence suggests an ancestral function in proteolysis that appeared 500 million years ago in the jawless vertebrates. In these organisms, the plasminogen gene is also present, but there are no plasminogen activators (1). Thus, ancient FSAP might have functioned as a primitive plasminogen activator, but the more recent, human-FSAP does not have this property (16). The EGF3 domain, which interacts with heparin, is strongly conserved across all vertebrates examined. In contrast, the histone interacting NTR is weakly conserved across vertebrates. Thus, the ability to detect and degrade histones might have evolved later and in some species only. FSAP has pro-coagulant, pro-fibrinolytic, histone-related- and cellular-functions that might have evolved along different paths with the changing requirements during evolution. In our analysis, did not consider other confounding factors such as the location of the gene in the genome, its expression pattern as well as its position in biological networks, which will all impact biological function. Although the activation of coagulation factors by charged surfaces is a well-established principle, the activation of FSAP zymogen with both positive and negative factors is unique, and its detailed analysis will help us to better understand zymogen activation in the context of vascular biology.

## Acknowledgements

We thank N. Bosvelt (Saxion University, Enschede, The Netherlands) for his excellent technical assistance and Dipankar Manna for the baculovirus expression trials: Funding was from the Research Council of Norway [Grant # 251239] to SMK and the Southeastern Norway Regional Health Authority (Grant 2015095) to BD. AC was supported by the Ministry of Education, Republic of Turkey (MoNE 1416/ YLSY). Calculations (or parts of them) for this publication were performed on the HPC cluster PALMA II of the University of Münster, subsidised by the DFG (INST 211/667-1). Andreas Lange was funded by the Volkswagen Stiftung (VWF), grant code 98183.

## Author contributions

SMK designed the study and supervised the experiments. SPSK constructed all the plasmids and performed all the expression and characterization studies and the AlphaFold analysis. AC and CK performed the evolution analysis. BD assisted with the analysis of the protein characterization. JE analyzed the docking studies and AL (Muenster) performed the molecular simulations. AL (Oslo) performed large-scale expression and purification. SMK drafted the manuscript. All authors added intellectual content to the manuscript as well as read, edited, and approved the final version of the manuscript.

## Data sharing statement

The data that support the findings of this study are available from the corresponding author upon reasonable request.

---

**Supplementary Fig. S1:**
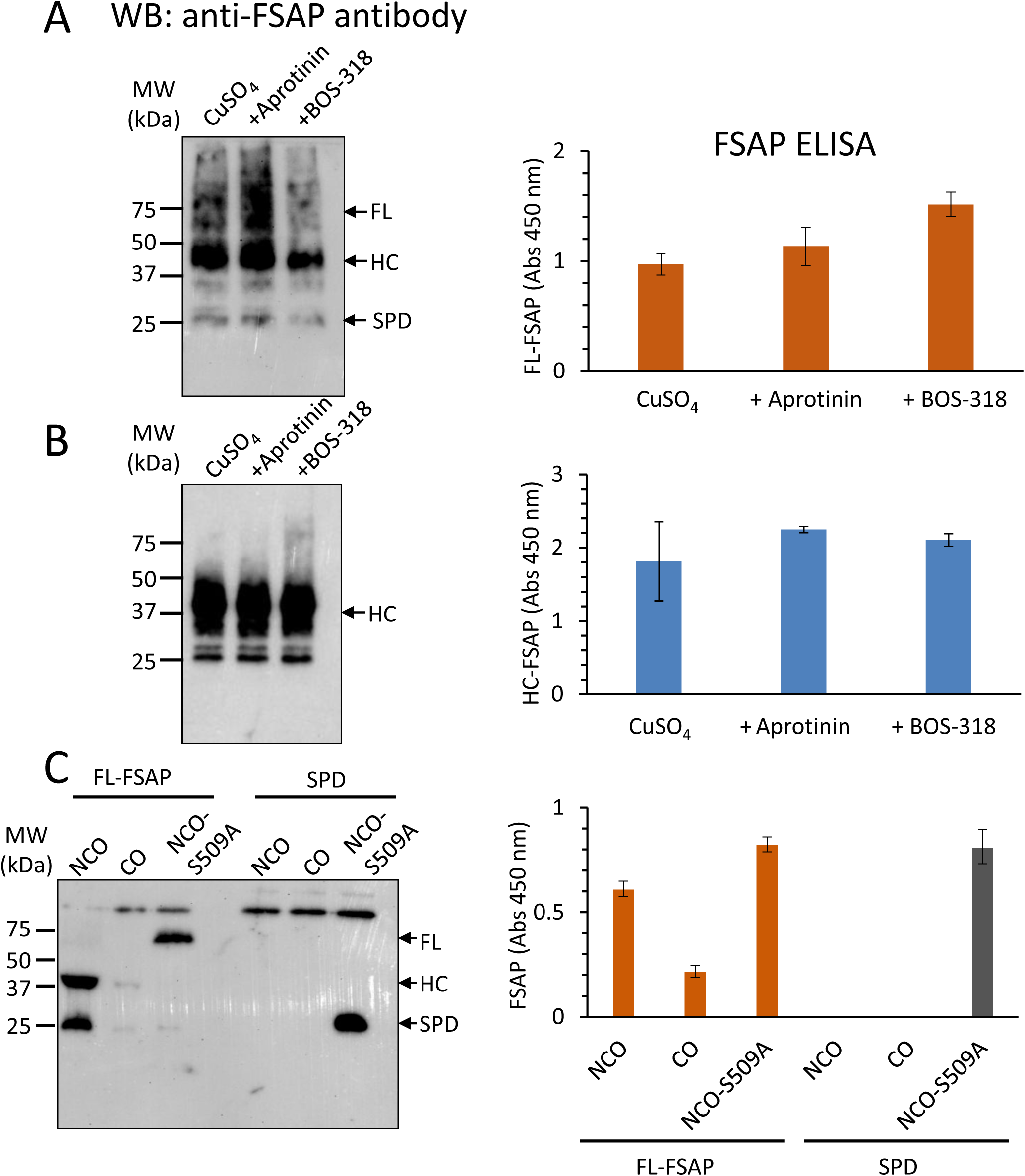
Effect of protease inhibition on recombinant FSAP expression: Effect of aprotinin (5 µg/ml) and BOS-318 (0.5 µM) on the expression of FL- **(A)** and HC-FSAP **(B)** in supernatants over 24 h. (**C)** Effect of mutating FSAP to inactivate the enzymatic activity (S509A). Left panels shows Western blot analysis with an anti-FSAP antibody and the right panel the results of an FSAP-ELISA. Similar results were obtained in three independent experiments. In the right panels, data is shown as mean ±SD (n=3 independent experiments).

**Supplementary Fig. S2:**
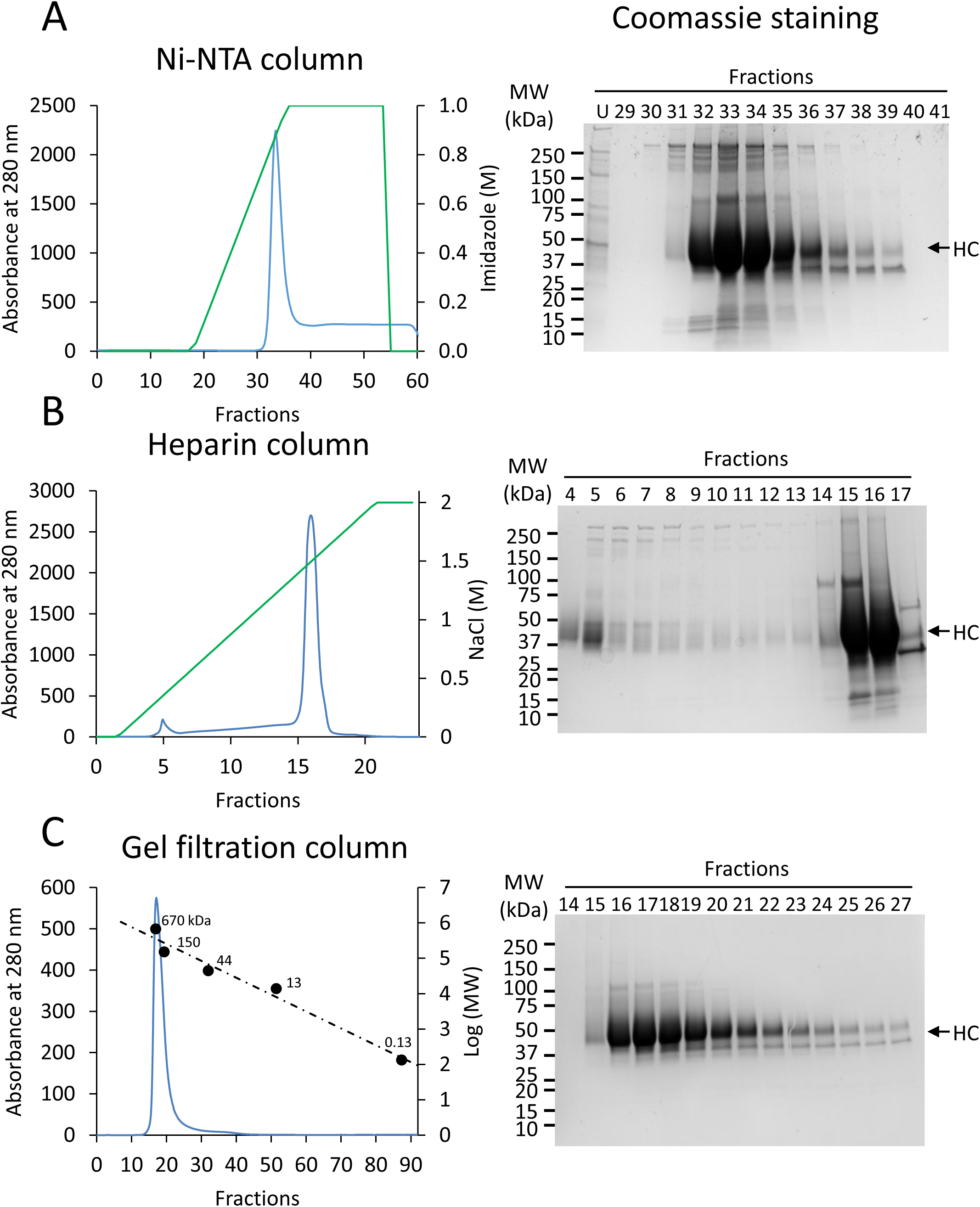
Purification of HC-FSAP: **A.** Ni-NTA chromatography. **B.** Heparin affinity chromatography. **C.** Size exclusion chromatography. Left panels show the chromatogram from each purification and right panels show SDS-PAGE analysis of the relevant fractions (arrow indicates position of HC-FSAP). Unbound (U) collected from Ni-NTA column; MW standards were used to calibrate the size exclusion column.

**Supplementary Fig. S3:**
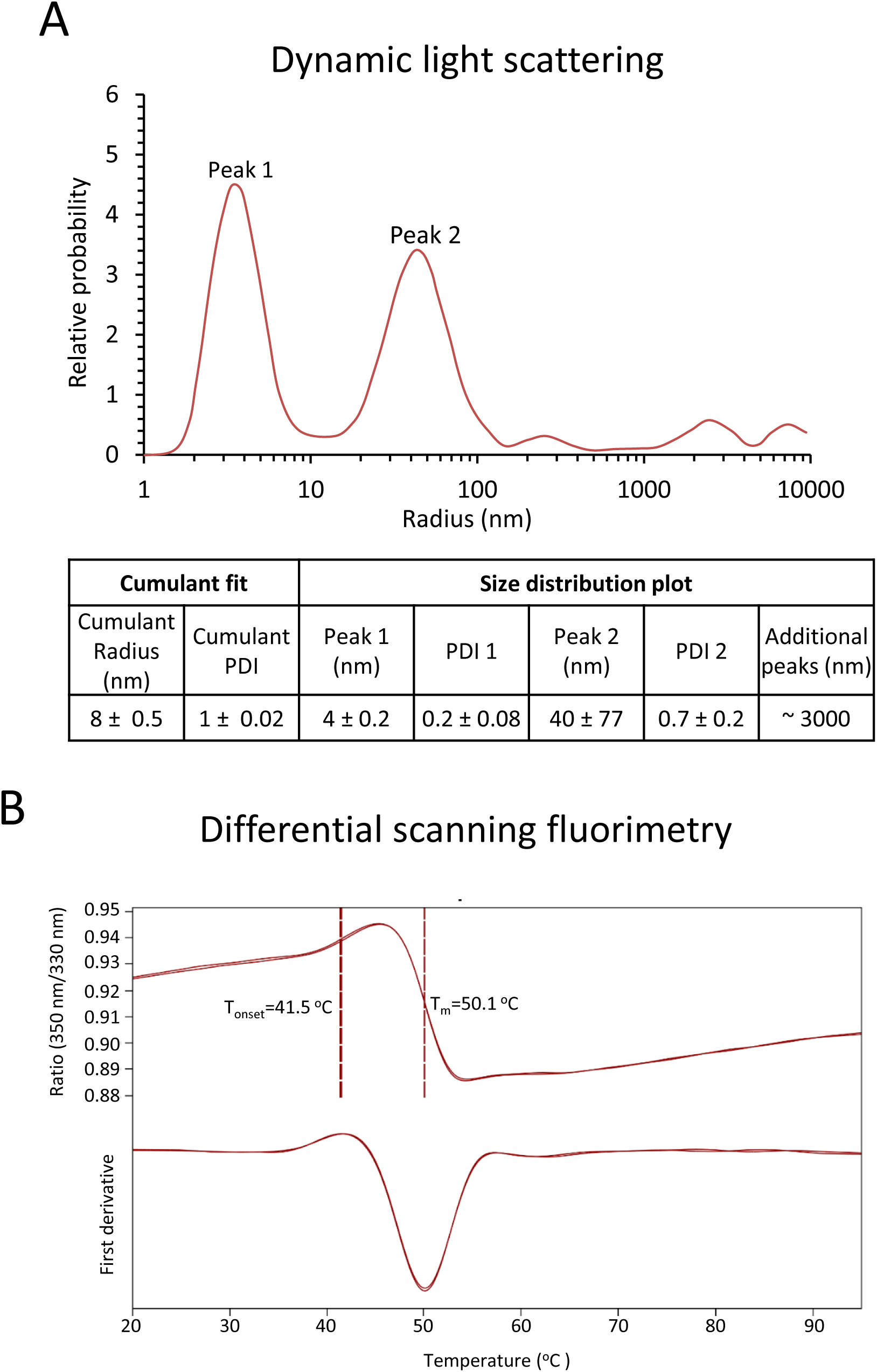
Biophysical characterization of purified HC-FSAP: **A.** Dynamic Light Scattering (DLS) measures the particle size distribution of HC-FSAP. The cumulant and size distribution fit of the particles is shown. **B.** Thermal stability and unfolding of HC-FSAP by nanoDSF (differential scanning fluorimetry) showing the fluorescence intensity ratio at 350 nm/330 nm (top) and the corresponding first derivative (bottom). The vertical lines depict the onset temperature (T_onset_) and melting temperature (T_m_).

**Supplementary Fig. S4:**
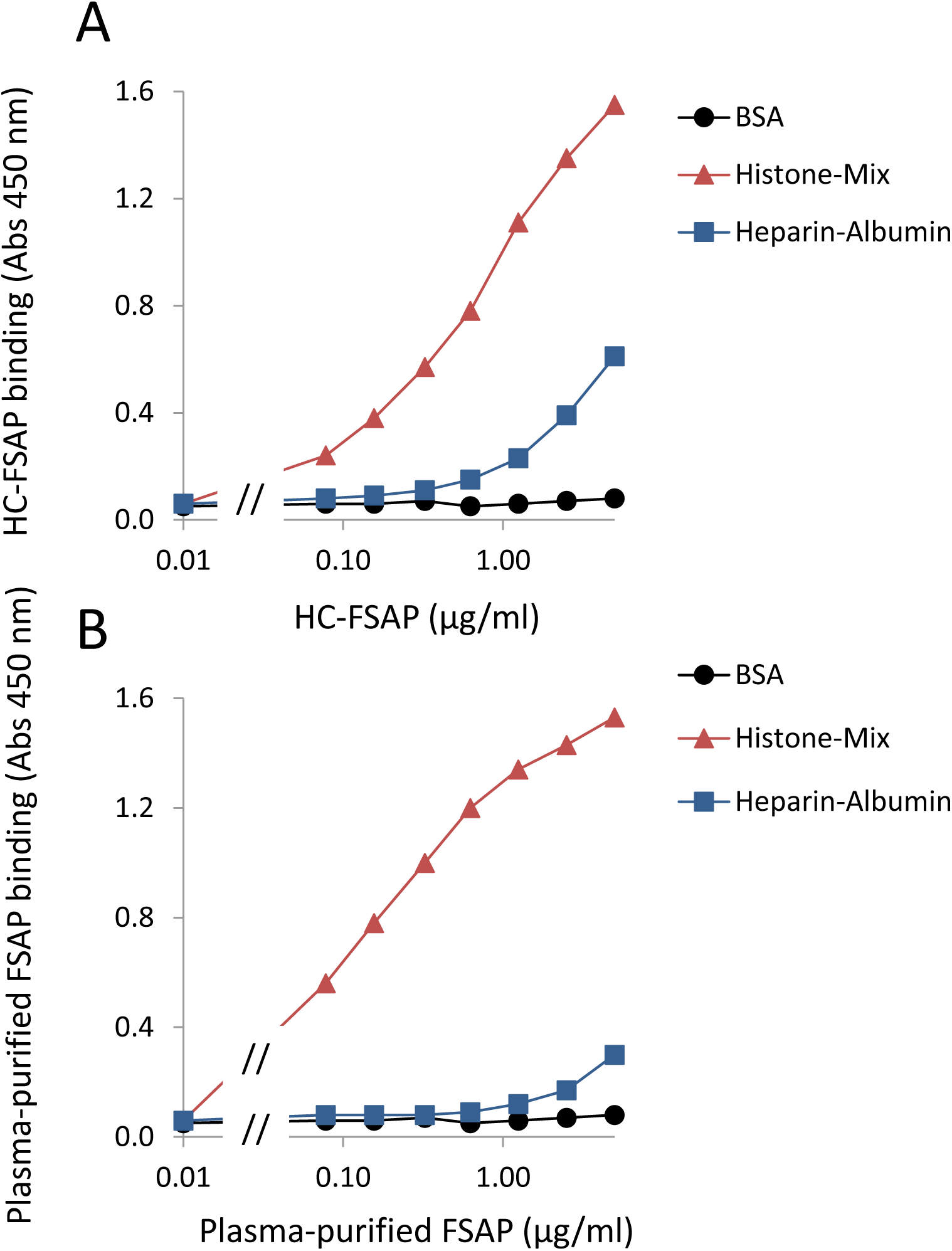
Binding of HC-FSAP and plasma-purified FSAP to Heparin and Histone: A mixture of bovine histones, heparin-albumin-biotin and bovine serum albumin (BSA) (all from Sigma) were coated (50 µl of 10 µg/ml solution) in wells of ELISA plates in coating buffer (pH 9.6). After blocking with BSA, either HC-FSAP **(A)** or plasma-purified FSAP **(B)** was added at a concentration of 0-5 µg/ml. Binding of FSAP was detected using an anti-FSAP antibody (mouse monoclonal) followed by an anti-mouse-peroxidase conjugate. Absorbance was measured after adding a colorimetric substrate. Data is shown as mean ±SD (n=3 replicates from one experiment). Error bars are smaller than the size of the symbols. Similar results were obtained in three independent experiments.

**Supplementary Fig. S5:**
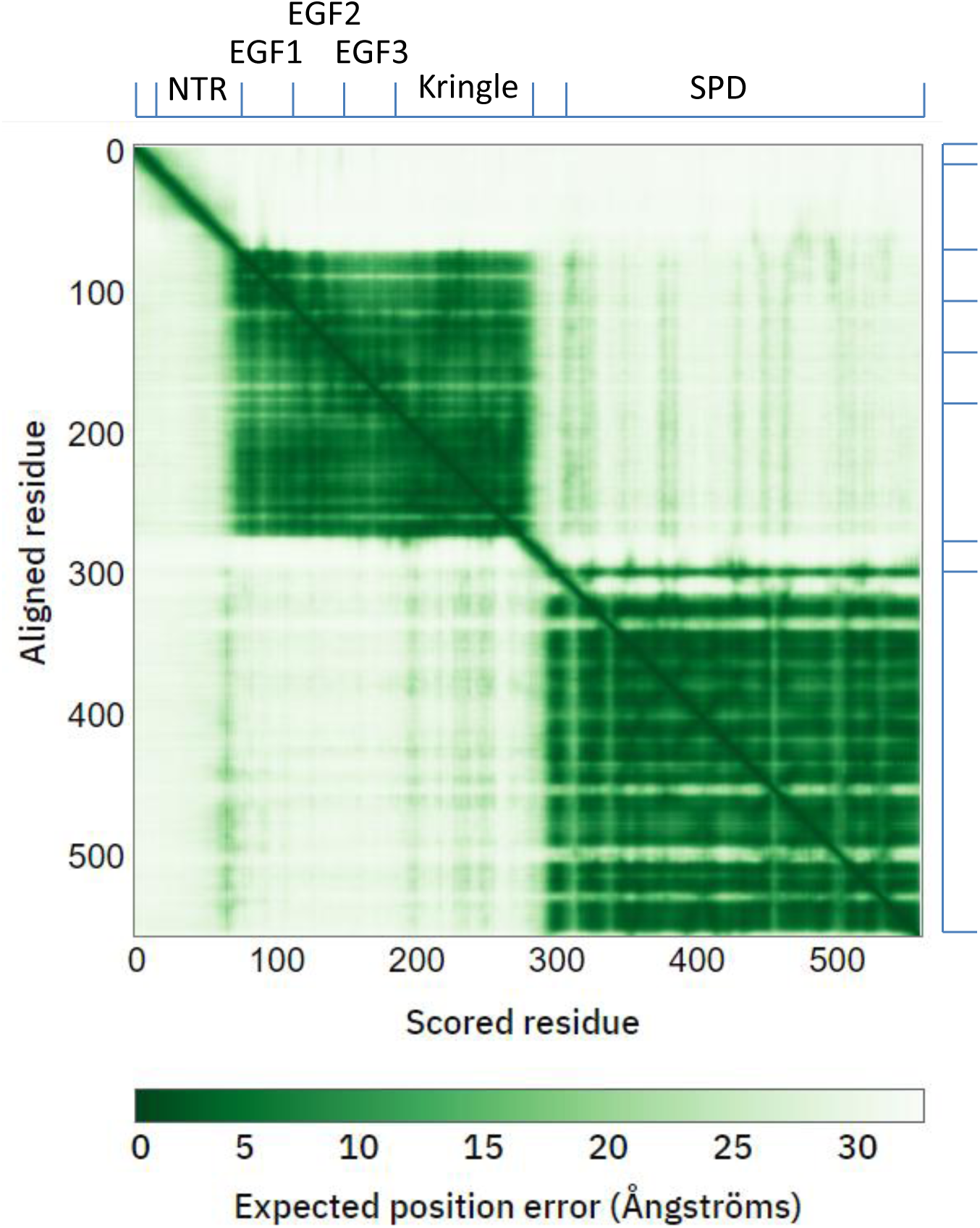
The predicted alignment error plot (PAE) plot: The AlphaFold-generated PAE plot of (FSAP: 1-560) is used to assess the relative domain position when residue Y is aligned to the expected position error on residue X. Interactions within the serine protease domain, EGF domains and kringle domains are likely to be accurate (dark green). Inter-domain interactions are likely to have high errors (light green to white).

**Fig. S6:**
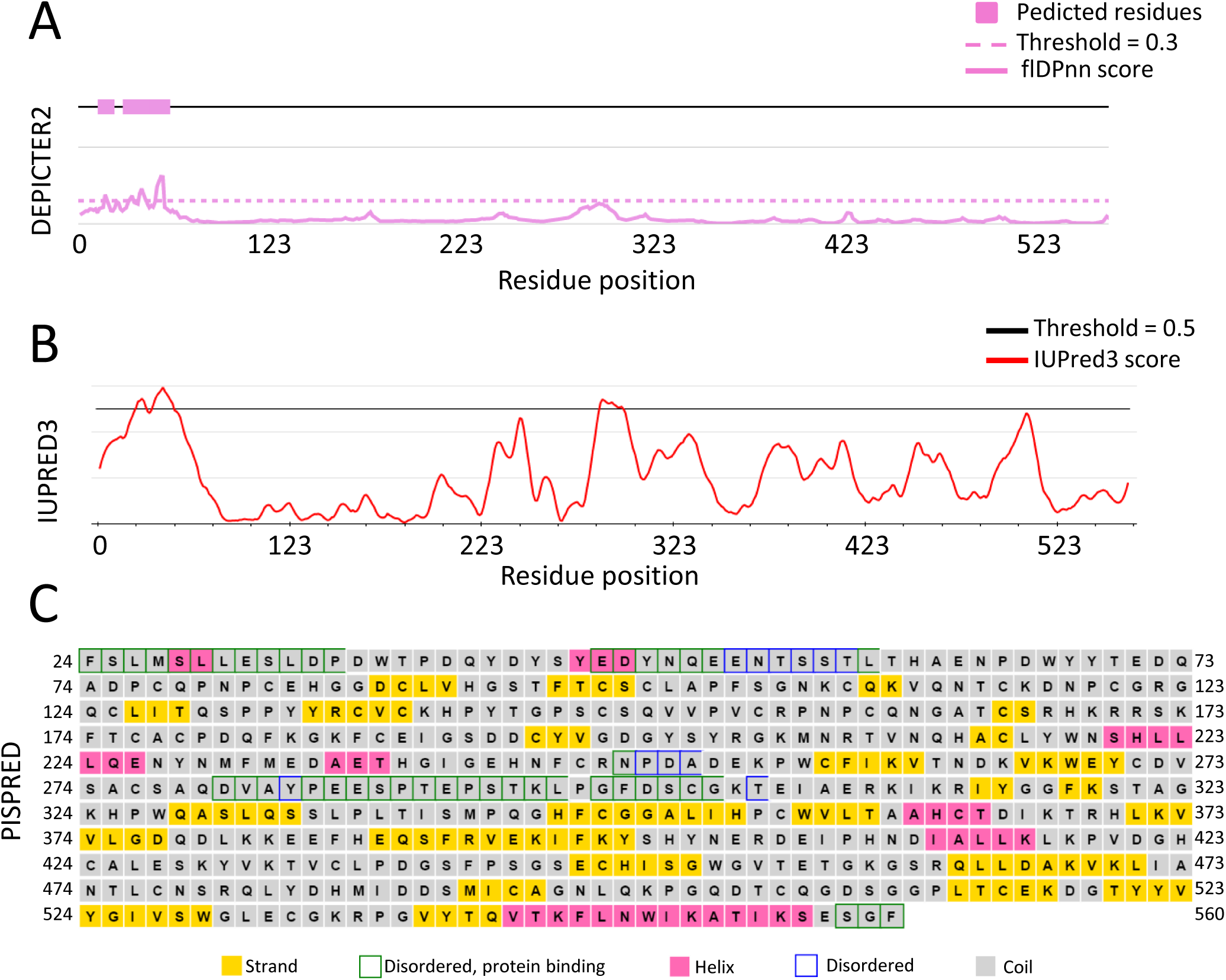
Intrinsic disorder prediction of FL-FSAP: All three programmes, DEPICTER2 **(A)**, IUPRED3 **(B)**, and PSIPRED **(C)** (FSAP:24-560) predict disorder in residues 50-60 of NTR, while PSIPRED also indicates disorder in the central and the C-terminal region.

**Supplementary Fig. S7.**
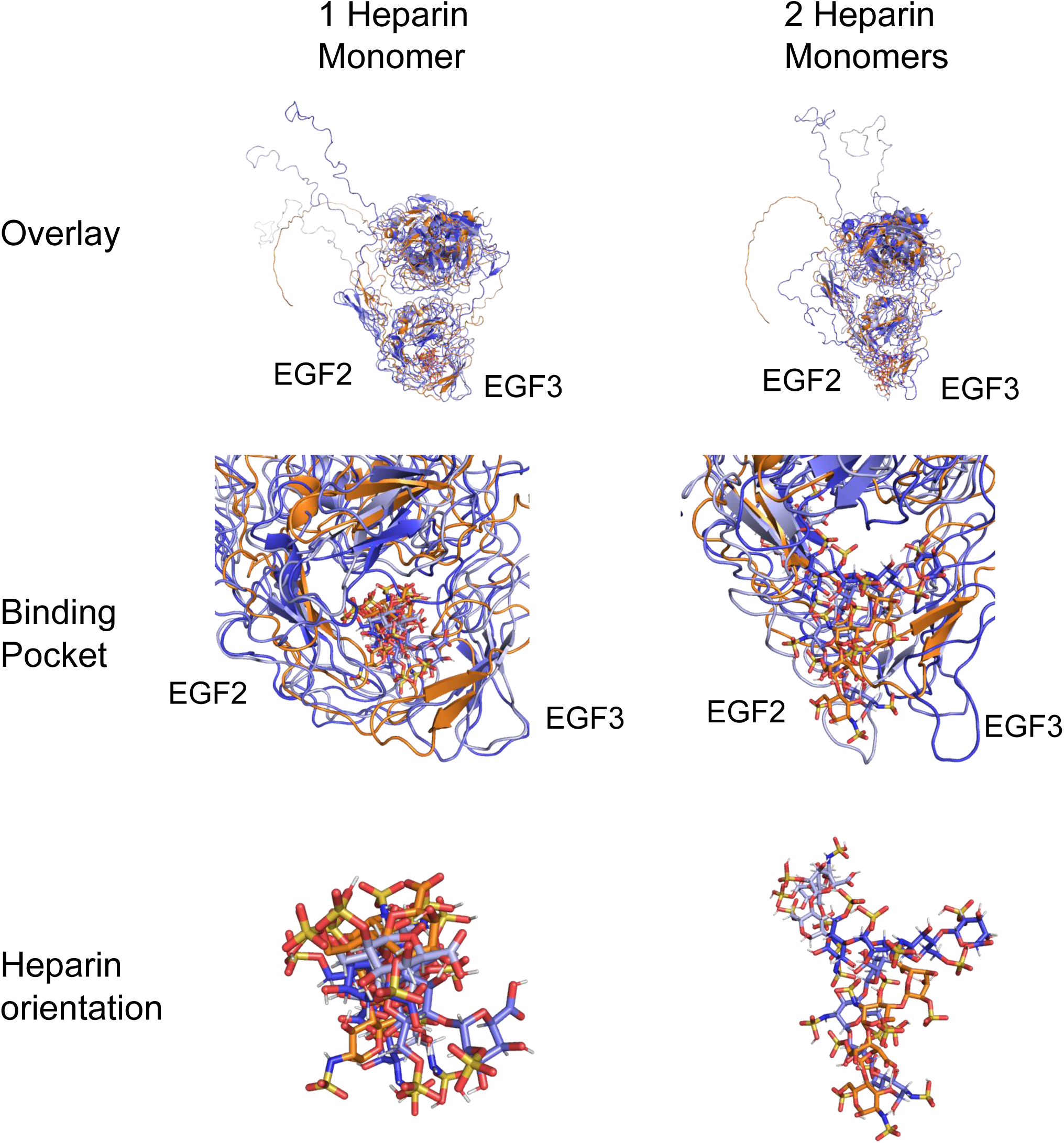
MD simulation of the FSAP protein with heparin as a ligand: examination of each heparin monomer separately: The first row presents an overlay of the starting structure and the three simulation outcomes. The initial structure is shown in orange, with simulation results after 200 ns displayed in light-blue (run1), slate-blue (run2), and dark blue (run3). The middle row focuses on the binding pocket, illustrating how the heparin monomers consistently remains within the pocket across all runs (color coding is the same as in row one). The final row shows the structure of the heparin monomers (each is a tetra-saccharide) after 200 ns in each simulation.

**Fig S8:**
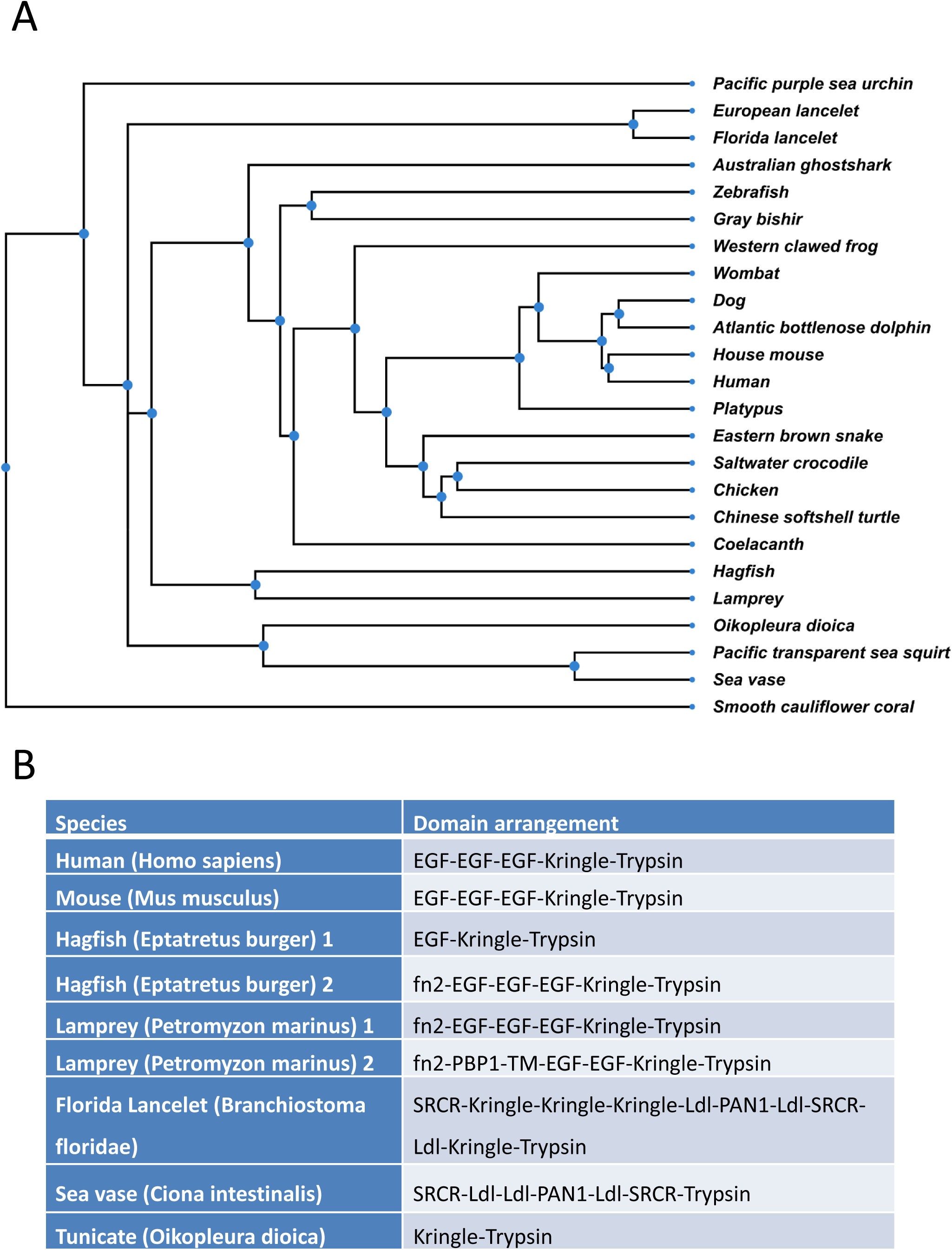
**A**. Phylogenetics tree and **B**. domain arrangement of putative orthologues of FSAP/HABP2 in the 24 selected species.

**Fig S9:**
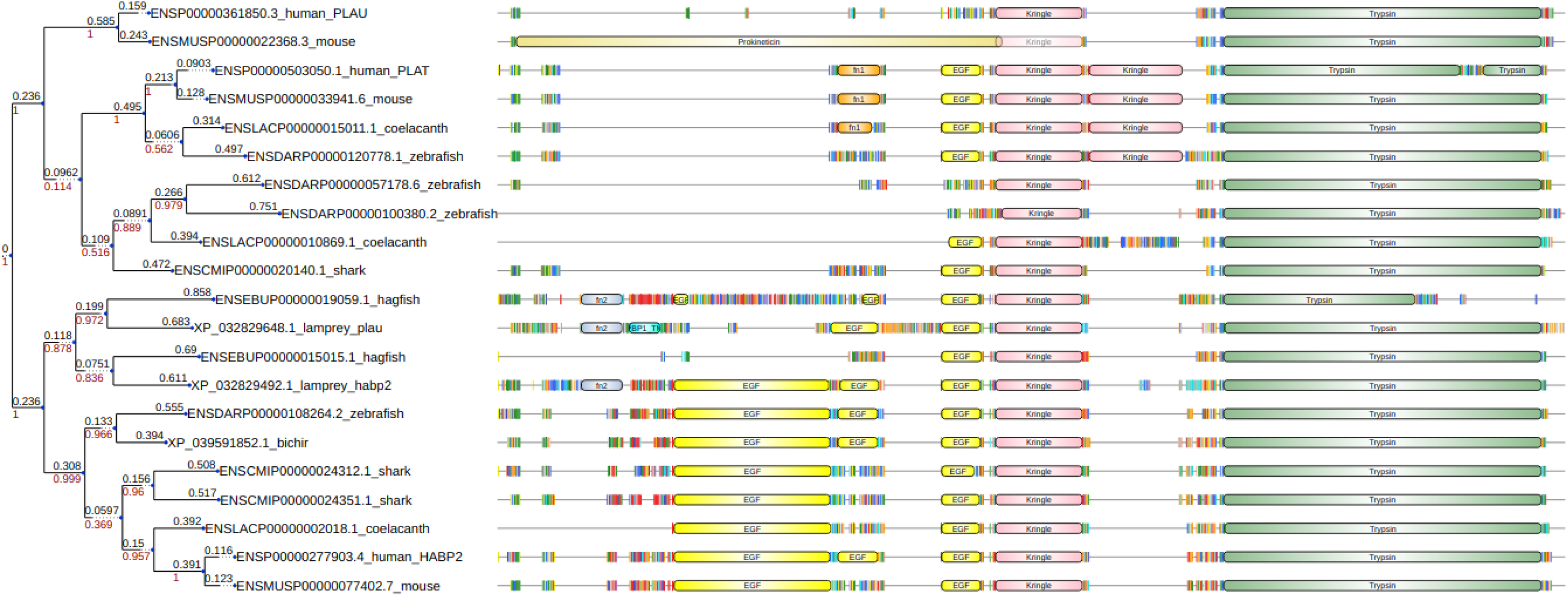
Phylogenetic tree of HABP2, PLAU and PLAT gene-encoded proteins (FSAP, uPA, tPA respectively) in vertebrates with their domain arrangement: Branch lengths are shown in black, bootstrap values in red. The alignment of the protein sequences is shown on the right side. Amino acids are indicated as colored vertical lines, domains are shown as boxes.

**Fig. S10:**
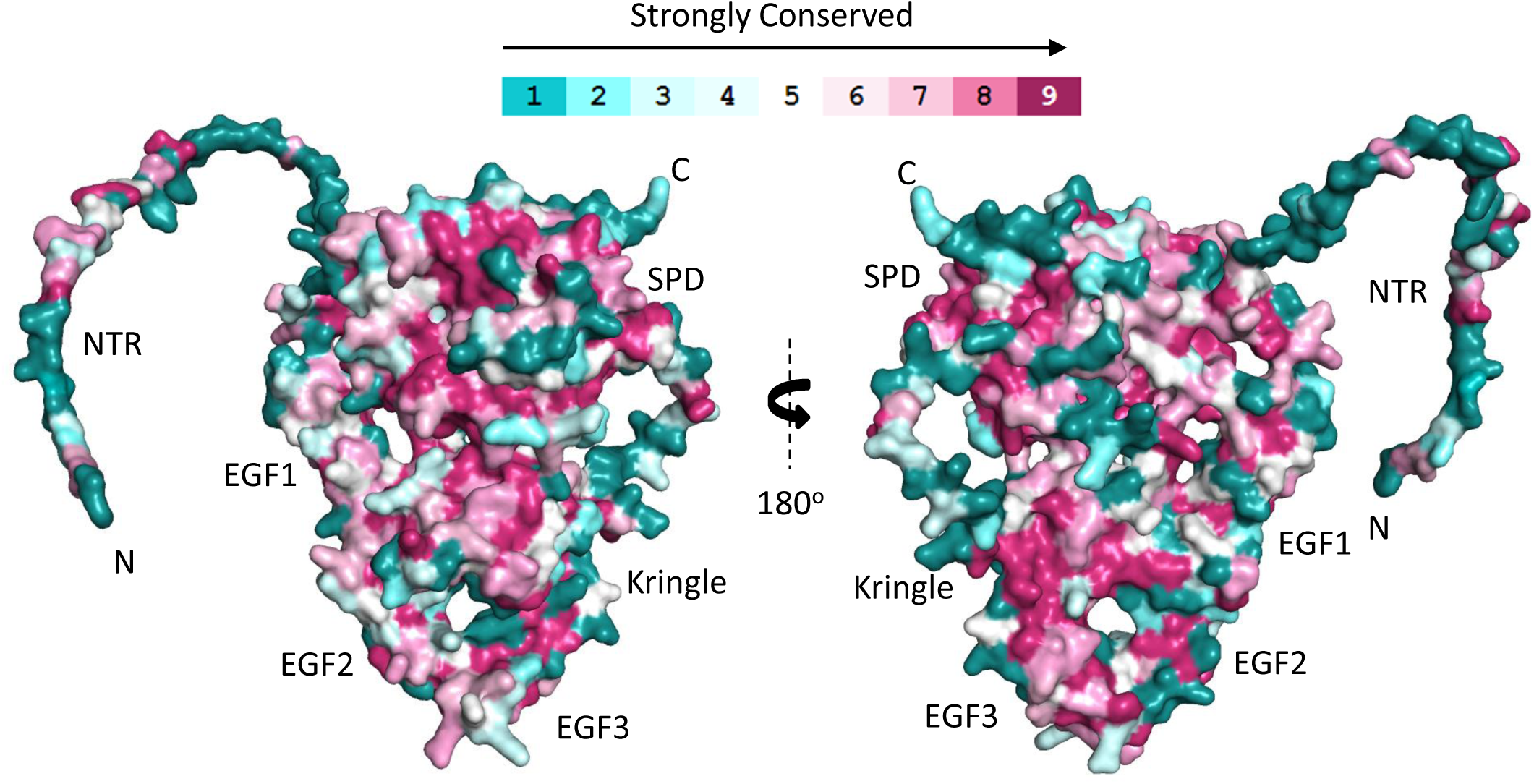
Conservation of full-length FSAP: FSAP sequence form 81 different species of the subphylum vertebrates were aligned using ConSurf. Maroon indicates high conservation and green indicates high variability.

